# N6-methyladenosine RNA methylation is a novel epitranscriptomic regulator of excessive alcohol drinking and vulnerability to relapse

**DOI:** 10.1101/2025.11.16.688690

**Authors:** Riccardo Maccioni, Irene Lorrai, Itzamar Torres, Roberta Puliga, Vez Repunte-Canonigo, Pietro Paolo Sanna

## Abstract

While internal RNA modifications have been known for decades, the contribution of epitranscriptomics to alcohol use disorder (AUD) remains unexplored. Here we investigated, for the first time, the role of the most abundant RNA modification, N6-methyladenosine (m^6^A) in regulating alcohol drinking and alcohol-induced gene expression. Mice with selective neuronal deletion of the m^6^A demethylase fat mass and obesity associated gene (*Fto*) showed enhanced initial motivation for alcohol, achieved escalated drinking more rapidly, and displayed greater relapse-like drinking after abstinence. Alcohol-naive *Fto*-deficient mice also exhibited potentiated alcohol-induced anxiolysis and sedation and blunted anxiogenic responses, despite unaltered alcohol metabolism. We then performed RNA enrichment coupled with RNA sequencing to characterize alcohol-induced epitranscriptomic and transcriptional remodeling. We observed that a history of alcohol exposure induced robust m^6^A hypermethylation in the hippocampus and that *Fto*-deficiency markedly altered m^6^A methylation and transcriptional dynamics. Gene Set Enrichment Analysis (GSEA) indicated that neuronal *Fto*-deficiency engages addiction-relevant pathways and that transcriptional programs associated with neuronal *Fto* deletion overlapped with alcohol-induced gene expression signatures, consistent with increased alcohol vulnerability. Our findings demonstrate that neuronal m^6^A RNA methylation is a novel regulator of excessive alcohol drinking and alcohol-dependent gene expression, and suggest that dysregulated epitranscriptomic control may contribute to the pathogenesis of AUD.

## Introduction

More than 170 distinct chemical modifications have been identified across coding and non-coding mammalian RNAs comprising the “epitranscriptome” ^1^ and contributing to the regulation of RNA expression and metabolism ^2–4^. Akin to epigenetic regulation of DNA, these internal RNA modifications influence its transcriptional efficiency, altering gene expression without changing the underlying genetic code ^3^. Among them, N6-methyladenosine (m^6^A) mRNA methylation is the most abundant and conserved modification ^5,6^. m^6^A mRNA modifications are preferentially deposited within the conserved motif DRACH (D = A/G/U, R = A/G, H = A/C/U), predominantly in 3’-UTRs and near stop codons, where they regulate splicing, nuclear export, stability, and translation ^7–9^. Multiple proteins are involved in the regulation of m^6^A RNA methylation, commonly referred as writers, erasers and readers that dynamically regulate m^6^A methylation and its functional output ^10^. Among these proteins, the fat mass and obesity associated protein FTO plays a key role in erasing m^6^A RNA methylation marks ^11^. While m^6^A has been found in mRNAs from diverse tissue types, the brain has especially high levels of m^6^A ^12^. RNA m^6^A methylation in the brain has been shown to modulate neuronal differentiation, synaptic plasticity, memory stabilization, and reward circuitry, and its dysregulation contributes to psychiatric disorders ^12–15^.

Here we demonstrate that neuronal *Fto* deficiency enhances the initial motivation for alcohol, accelerates the development of alcohol dependence, and increases relapse-like drinking during withdrawal. A history of dependent alcohol exposure elevated m^6^A methylation in the hippocampus during acute withdrawal. We also observed that neuronal *Fto* deletion broadly altered m^6^A RNA methylation and gene expression in both alcohol-naive and alcohol-dependent mice, affecting genes involved in synaptic dynamics, monoaminergic and GABAergic neurotransmission, and addiction-related signaling pathways. Notably, *Fto* deficiency partially recapitulated alcohol-induced transcriptional programs. Collectively, these findings identify FTO-mediated m^6^A regulation as a previously unrecognized contributor to excessive alcohol drinking and vulnerability to alcohol dependence.

## Results

### Deletion of neuronal *Fto* accelerates acquisition of alcohol drinking, onset of alcohol dependence and promotes relapse-like behavior

We bred *Fto* floxed mice (*Fto-fl/fl*) with the Synapsin-1-Cre deleter line to obtain mice with homozygous neuron-specific deletion of *Fto* (henceforth, *Fto-Syn1-Cre* mice). We first measured initial preference for alcohol, followed by 4 weeks of non-dependent alcohol-drinking using a two-bottle choice (2BC) paradigm (Fig.1a). Alcohol-naive *Fto-Syn1-Cre* mice consumed significantly more alcohol than control *Fto-fl/fl* mice during their first alcohol exposure (Fig.1b). However, average alcohol intake did not differ between the two genotypes over 4 weeks (Fig.1c).

**Fig. 1.**
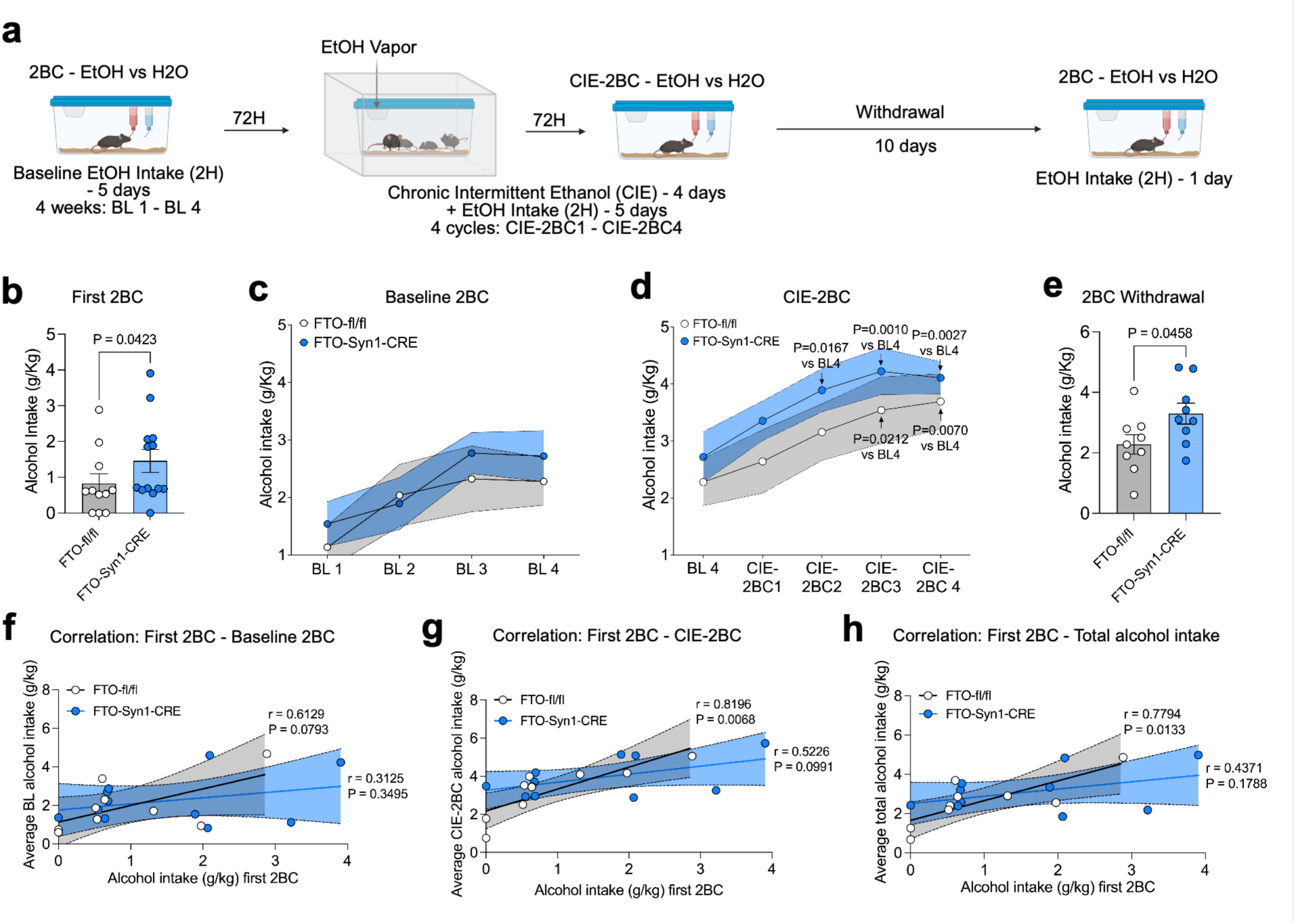
Genetic deletion of *Fto* in neurons enhanced initial motivation for alcohol, hastened alcohol dependence, and increased relapse-like drinking. a) Timeline of experimental procedures. **b)** Alcohol intake (g/kg) measured after the first two-bottle choice (2BC) in *Fto-fl/fl* (n=11) and *Fto-Syn1-Cre* (n=13) mice. Statistical significance was determined by two-tailed Mann-Whitney test. **c)** Average weekly alcohol intake (g/kg) of 4 weeks of baseline measurements (BL1 - 4) in *Fto-fl/fl* (n=9) and *Fto-Syn1-Cre* (n=11) mice. **d)** Alcohol intake (g/kg) during CIE-2BC in *Fto-fl/fl* (n=9) and *Fto-Syn1-Cre* (n=11) mice. Statistical significance was determined by two-way repeated measure (RM) ANOVA followed by Tukey’s multiple comparisons test. **e)** Alcohol intake (g/kg) during the single 2BC conducted after 10 days of withdrawal in *Fto-fl/fl* (n=9) and *Fto-Syn1-Cre* (n=9) mice. Statistical significance was determined by two-tailed Welch’s t test. Pearson’s correlation analysis between the initial intake of alcohol measured during the first 2BC and the average of non-dependent (BL 1-4) alcohol intake **(f)**, the average alcohol consumption during the CIE-2BC cycles **(g)** and the average of total alcohol consumption along the whole procedure **(h)**.

To assess the role of neuronal *Fto* in dependence-associated drinking, we induced alcohol dependence by repeated cycles of alcohol vapor exposure with the Chronic Intermittent Ethanol paradigm coupled with 2 Bottle Choice (CIE-2BC). *Fto-Syn1-Cre* mice only required 2 cycles of CIE-2BC to achieve significantly escalated (dependent) alcohol drinking, while 3 cycles were required for *Fto-fl/fl* mice (Fig.1d). Blood alcohol levels (BALs) did not differ between *Fto-Syn1-Cre* and *Fto-fl/fl* mice after vapor exposure, indicating that the two genotypes were exposed to equal amounts of alcohol and that neuronal *Fto* deletion does not alter alcohol metabolism (Supplementary Figure S1, Supplementary Information).

We then tested the role of neuronal *Fto* in reinstatement behavior. To this end, withdrawal was induced in dependent *Fto-Syn1-Cre* and *Fto-fl/fl* mice by preventing alcohol access for 10 days (Fig.1a). When mice were re-exposed to alcohol in the 2BC, *Fto-Syn1-Cre* mice consumed a significantly greater amount of alcohol than *Fto-fl/fl* control mice (Fig.1e).

Finally, we analyzed the correlation between initial alcohol intake during the first 2BC session (alcohol-naive) and later (baseline, CIE-2BC and total) consumption of alcohol in the two genotypes. While in *Fto-fl/fl* mice initial drinking correlated with subsequent drinking, no such association was observed in *Fto-Syn1-Cre* mice (Fig.1f-h).

Collectively, these results indicate that neuronal *Fto* deletion alters alcohol-drinking trajectories and promotes a more vulnerable phenotype characterized by enhanced early alcohol motivation, accelerated development of dependence, and increased relapse-like behavior.

### Deletion of neuronal *Fto* potentiates alcohol-induced anxiolysis and prevents alcohol-induced anxiogenic response

We then evaluated the contribution of neuronal *Fto* to acute alcohol-induced anxiety-like behavior in the Elevated Plus Maze (EPM) and Light and Dark Transition (LDT) tests (Fig.2a). In the EPM, acute alcohol administration produced a stronger anxiolytic effect in *Fto-Syn1-Cre* mice, as indicated by a significantly higher percentage of time spent in the open arms compared with alcohol-treated *Fto-fl/fl* controls (Fig.2b). As expected, alcohol produced significant anxiolytic effects in both *Fto-Syn1-Cre* and *Fto-fl/fl* mice in both time spent and entries into the open arms compared to saline-administered mice (Fig.2b,c). No significant differences between *Fto-fl/fl* and *Fto-Syn1-Cre* mice were observed upon saline administration for either parameter (Fig.2b,c). Importantly, measures of total entries in both open and closed arms excluded alcohol-induced sedation in either genotype and indicated a higher level of activity in *Fto-Syn1-Cre* than *Fto-fl/fl mice* (Fig.2d).

**Fig. 2.**
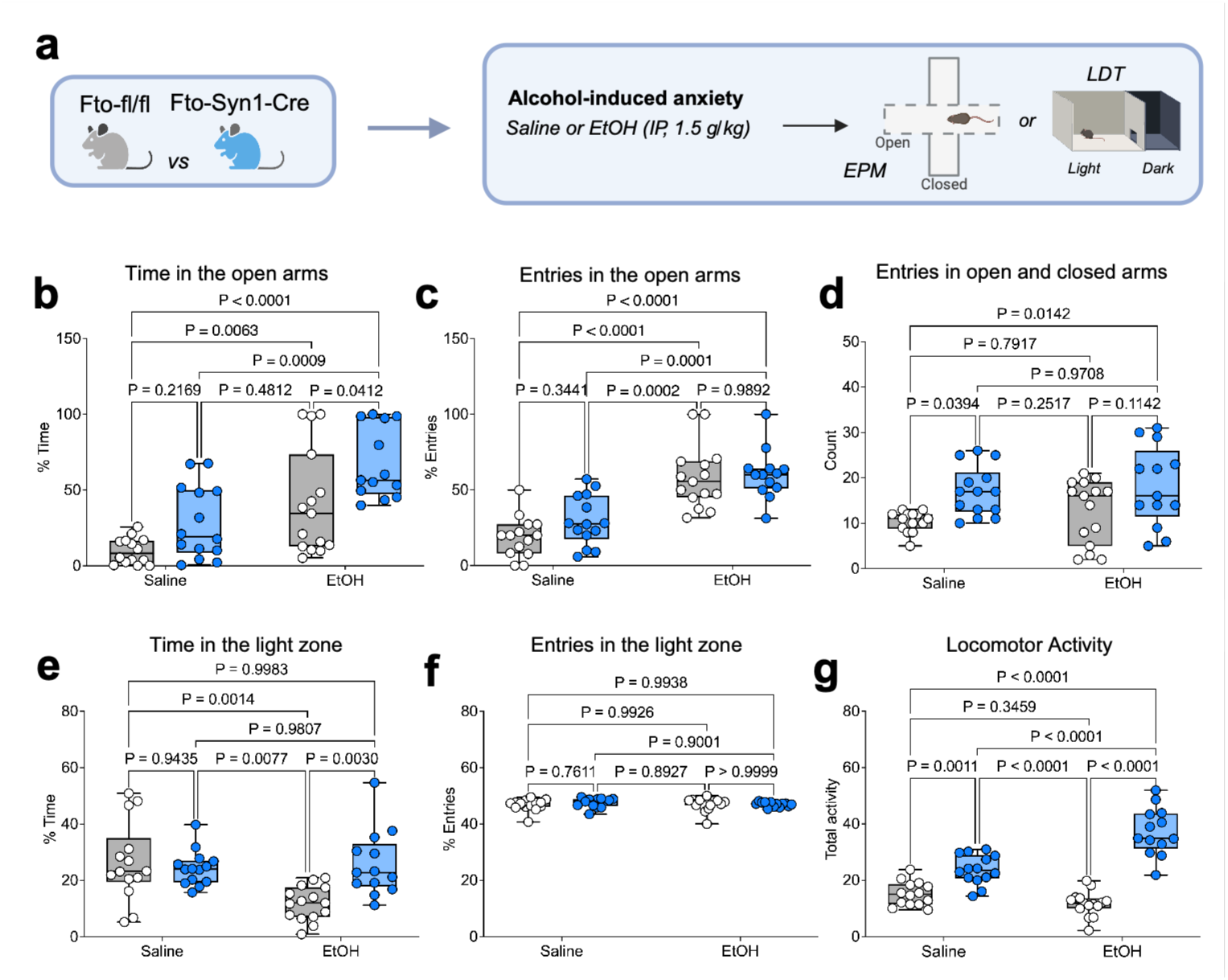
Genetic deletion of *Fto* in neurons increases alcohol-induced anxiolysis and anxiogenic responses. **a)** *Fto-fl/fl* and *Fto-Syn1-Cre* mice were assessed at the Elevated Plus Maze (EPM) and the Light and Dark Transition (LTD) test after 5 minutes from a single acute IP administration of alcohol (1.5 g/kg). Percentage of time **(b)** and entries **(c)** in the open arms of *Fto-fl/fl* and *Fto-Syn1-Cre* mice with and without alcohol treatment. **d)** Total entries in the open arms of *Fto-fl/fl* and *Fto-Syn1-Cre* mice. Percentage of time **(e)** and entries **(f)** in the Light Zone (LZ) of *Fto-fl/fl* and *Fto-Syn1-Cre* mice with and without alcohol treatment. **g)** Total locomotor activity of *Fto-fl/fl* and *Fto-Syn1-Cre* mice. Statistical significance was determined by two-way ANOVA followed by Tukey’s multiple comparisons test (n=13-15).

Conversely, alcohol induced anxiogenic-like response in the LDT in *Fto-fl/fl* mice, as reflected by a significant reduction in time spent in the light zone. This effect was not observed in *Fto*-*Syn1-Cre* mice, in which alcohol administration did not reduce time in the light compartment and remained significantly higher than in alcohol-treated *Fto*-*fl/fl* mice (Fig.2e). Interestingly, we did not observe any difference in the percentage of entries in the light zone (Fig.2f). Similarly to what was observed in the EPM, no effect of genotype was detected in mice that were administered saline before the test (Fig.2e,f), while we observed a strong effect of genotype in mediating locomotor activity, with *Fto-Syn1-Cre* mice generally more active than *Fto-fl/fl* mice, in particular upon alcohol treatment (Fig.2g).

Collectively, these findings suggest that neuronal *Fto* deletion modulates behavioral sensitivity to the acute anxiolytic and anxiogenic effects of alcohol.

### Deletion of neuronal *Fto* potentiates alcohol-induced sedative and hypnotic effects

Since alcohol-induced anxiolysis and sedation share overlapping mechanisms ^16,17^, we also assessed *Fto-fl/fl* and *Fto-Syn1-Cre* mice in the loss of righting reflex (LORR) to evaluate alcohol-induced sedation and hypnosis (Fig.3a). Sedation was rapidly induced by an acute injection of an intoxicating dose of alcohol (3.5 g/kg) in both genotypes (Fig.3b). In keeping with the enhanced anxiolytic response to alcohol, *Fto-Syn1-Cre* mice also exhibited greater sensitivity to alcohol-induced hypnotic and sedative effects, as indicated by the significant increase in LORR duration (Fig.3c). Measurement of BALs from blood samples collected at awakening and 2 h after alcohol injection did not reveal any difference between *Fto-fl/fl* and *Fto-Syn1-Cre* mice, ruling out differences in alcohol metabolism between the genotypes (Fig.3d).

**Fig. 3.**
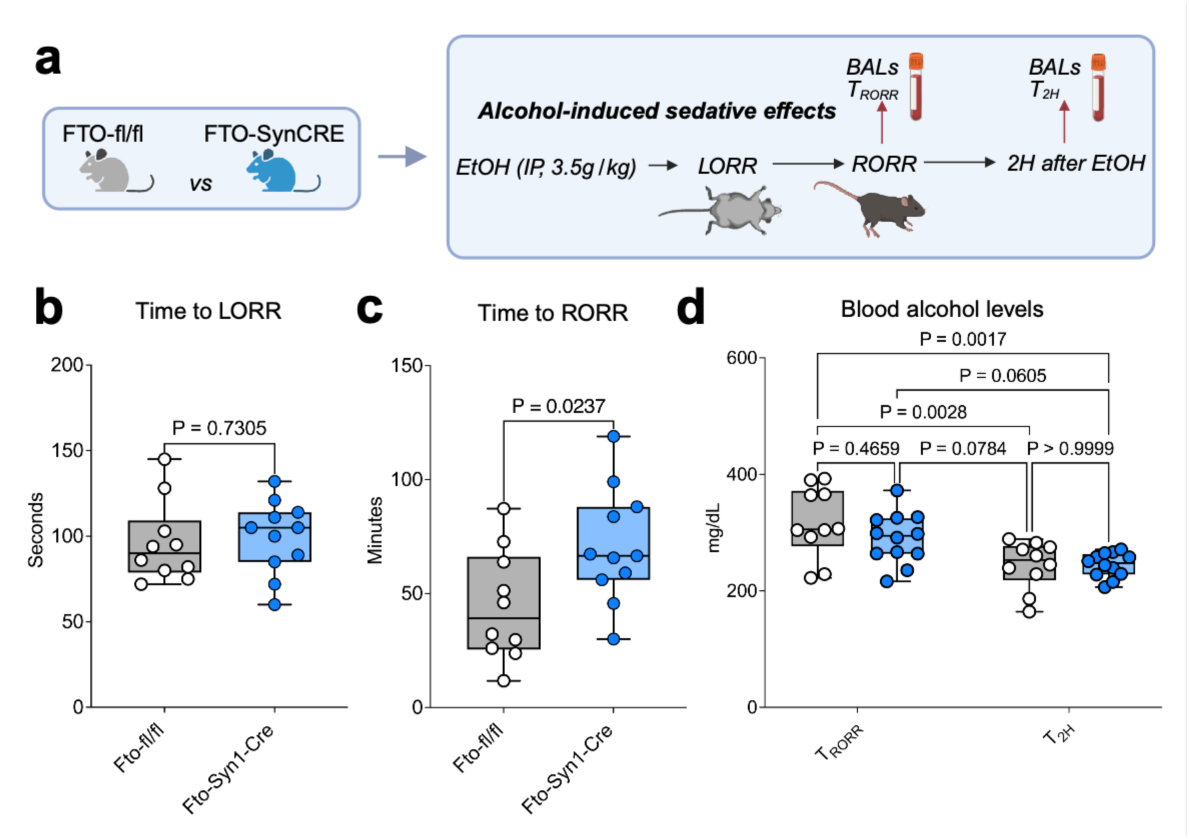
Increased sensitivity to the sedative effects of alcohol in mice with neuronal *Fto* deficiency. **a)** Alcohol (3.5 g/kg, IP) induced loss of Righting reflex (LORR) in *Fto-fl/fl* and *Fto-Syn1-Cre mice*. *Fto-Syn1-Cre* mice showed unchanged time to sedation **(b)** but higher duration of sedation as measured by the time to recover the righting reflex (RORR) **(c)**. Statistical significance was determined by two-tailed Welch’s t test (n=10-11). **(d)** No differences between the two genotypes were observed in the blood alcohol levels (BALs). Statistical significance was determined by two-way RM ANOVA followed by Tukey’s multiple comparisons test (n=10-11).

### Alcohol dependence increases RNA m^6^A methylation and differentially remodels neuronal and signaling pathways under neuronal *Fto* deletion

To investigate the effect of alcohol on m^6^A RNA methylation, we performed the EpiPlex^TM^ assay, an integrated assay and analysis platform for m^6^A and RNA expression profiling, using hippocampal RNA from alcohol-naive and alcohol-dependent *Fto-fl/fl* and *Fto-Syn1-Cre* mice after 24 hours of withdrawal. Gene Set Enrichment Analysis (GSEA) was then applied on our EpiPlex^TM^ data to characterize differentially m^6^A methylated genes. We observed no effect of genotype or alcohol on the proportion of m^6^A peaks overlapping with the canonical DRACH motif across conditions (Fig.4a) or on the genomic distribution of m^6^A peaks across transcript regions (Fig.4b). Principal Component Analysis (PCA) of m^6^A methylation profiles revealed clear separation across genotypes and treatment conditions, indicating robust effects of neuronal *Fto* deletion and alcohol treatment (Fig.4c).

**Fig. 4.**
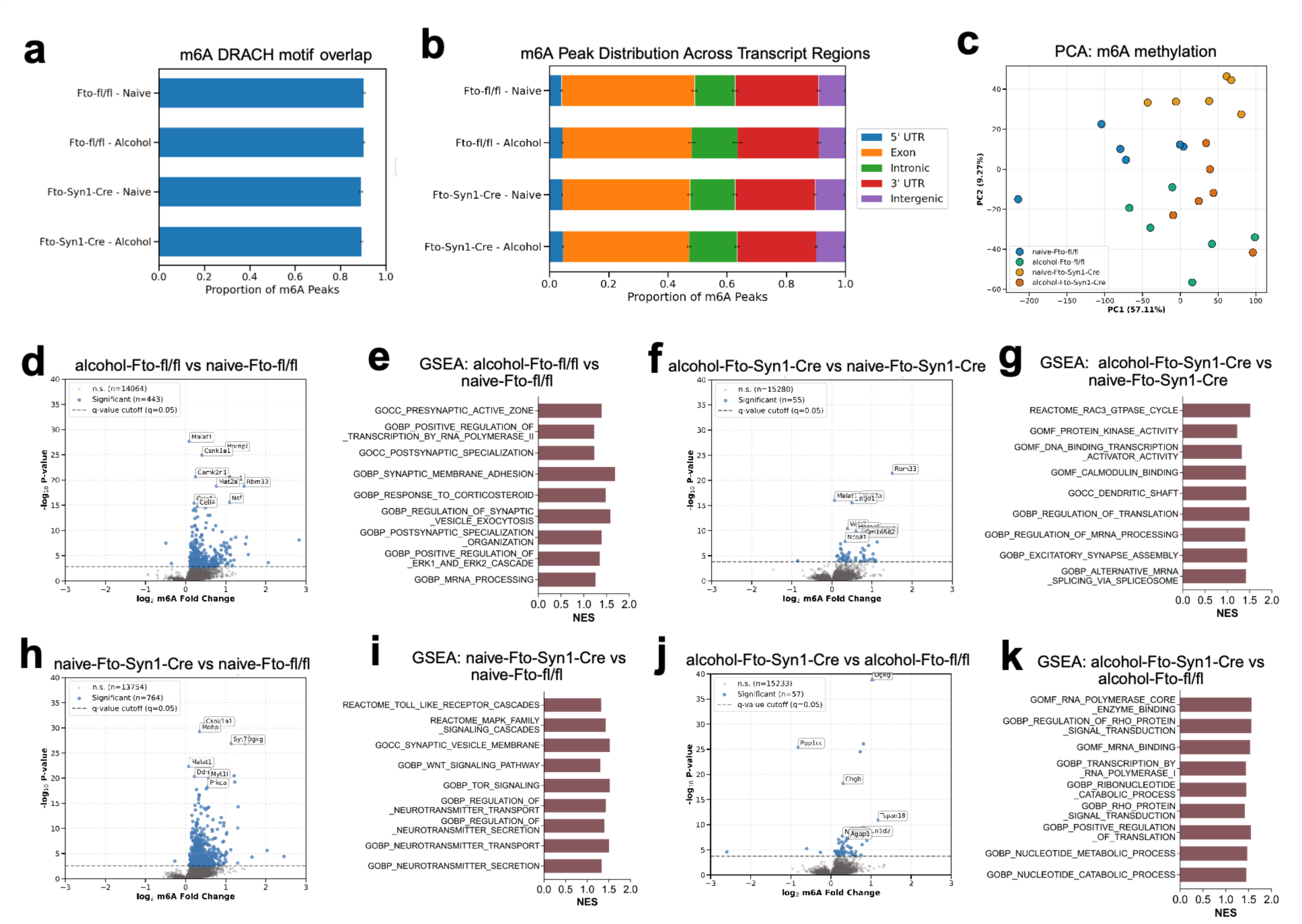
Alcohol dependence increases hippocampal m^6^A methylation and reveals *Fto*-dependent m^6^A epitranscriptomic remodeling. **a)** Proportion of m^6^A peaks overlapping with the canonical DRACH motif in hippocampal samples from alcohol-naive and alcohol-dependent *Fto-fl/fl* and *Fto-Syn1-Cre* mice. **b)** Genomic distribution of m^6^A peaks across transcript regions (5′UTR, exon, intron, 3′UTR, and intergenic regions). **c)** Principal Component Analysis (PCA) of enriched m^6^A modification signal detected by the EpiPlex™ assay binned by gene-body regions. Genotype and treatment replicates are indicated by color: alcohol-naive *Fto-fl/fl* (blue, n=6), alcohol-dependent *Fto-fl/fl* (green, n=6), alcohol-naive *Fto-Syn1-Cre* (orange, n=6), alcohol-dependent *Fto-Syn1-Cre* (red, n=6). **d, f, h, j)** Volcano plots of differentially m^6^A methylated peaks in alcohol-dependent *Fto-fl/fl* (vs alcohol-naive *Fto-fl/fl*) (d), alcohol-dependent *Fto-Syn1-Cre* (vs alcohol-naive *Fto-Syn1-Cre*) (f), alcohol-naive *Fto-Syn1-Cre* (vs alcohol-naive *Fto-fl/fl* (h) and alcohol-dependent *Fto-Syn1-Cre* (vs alcohol-dependent *Fto-fl/fl* (j). Transcript regions with significant increase or decrease in m^6^A methylation were identified by EpiScout™ software (FDR-corrected P-value < 0.05). The direction of the change is relative to the first term of the comparison. **e, g, i, k)** Representative pathways enrichment analysis. Bars represent NES (Normalized Enrichment Scores) of differentially m^6^A methylated transcripts. The direction of the change is relative to the first term of the comparison. Selected pathways with P-value < 0.05 are shown.

Alcohol-dependence induced a marked and predominantly unidirectional change in m^6^A methylation in control *Fto-fl/fl* mice, with 443 (441 hypermethylated and 2 hypomethylated) differentially modified m^6^A regions (DMRs) in alcohol-dependent compared to alcohol-naive littermates (Fig.4d). GSEA revealed enrichment of pathways involved in synaptic structure and vesicle dynamics, ERK-dependent signaling, transcriptional regulation, and mRNA processing. Gene sets related to corticosteroid response were also enriched (Fig.4e).

In contrast, the comparison between alcohol-dependent *Fto-Syn1-Cre* and alcohol-naive *Fto-Syn1-Cre* mice yielded a considerably smaller number of DMRs, with 55 (54 hypermethylated, 1 hypomethylated) DMRs found in alcohol-dependent *Fto-Syn1-Cre* mice (Fig.4f). GSEA indicated enrichment of pathways related to synaptic structural remodeling and activity-dependent signaling, including Rac GTPase cycling, protein kinase activity, calmodulin binding, and excitatory synapse assembly. In parallel, gene sets involved in regulation of translation, mRNA processing, and alternative splicing were also enriched (Fig.4g).

The reduced number of alcohol-induced DMRs in *Fto-Syn1-Cre* mice may reflect their elevated m^6^A methylation baseline. Accordingly, 764 (763 hypermethylated, 1 hypomethylated) DMRs were found in alcohol-naive *Fto-Syn1-Cre* when compared to alcohol-naive *Fto-fl/fl* (Fig.4h). GSEA revealed enrichment of pathways related to synaptic signaling and intracellular signaling cascades, including neurotransmitter transport and secretion, synaptic vesicle membrane components, MAPK family signaling, WNT signaling, TOR signaling, and Toll-like receptor cascades (Fig.4i).

Finally, 57 (53 hypermethylated, 4 hypomethylated) DMRs were found in alcohol-dependent *Fto-Syn1-Cre* mice when compared to alcohol-dependent *Fto-fl/fl* (Fig. 4j). Enriched pathways included transcriptional regulation and RNA metabolism, including RNA polymerase core enzyme binding, transcription by RNA polymerase II, mRNA binding, ribonucleotide catabolic processes, and nucleotide metabolic pathways. Gene sets associated with Rho protein signal transduction and positive regulation of translation were also enriched (Fig.4k).

Collectively, these results reveal that alcohol dependence drives robust and predominantly hypermethylating changes in the hippocampal m⁶A landscape, an effect that is substantially modified by neuronal *Fto* deletion. All differentially methylated transcripts and pathways can be found in the Supplementary Tables (Supplementary Table 1, Supplementary Table 2, Supplementary Table 3, Supplementary Table 4, Supplementary Table 5, Supplementary Table 6, Supplementary Table 7, Supplementary Table 8). Quality control (QC) can be found in the Supplementary Information (Supplementary Figure S2).

### Neuronal *Fto* deletion converges with and amplifies alcohol-induced transcriptional remodeling of synaptic and signaling pathways

RNA-Seq analysis was also carried out using the EpiPlex^TM^ assay from the hippocampus of alcohol-naive and alcohol-dependent *Fto-fl/fl* and *Fto-Syn1-Cre* mice after 24 hours of withdrawal. GSEA was then applied to our RNA-seq data to identify molecular pathways transcriptionally regulated in an *Fto*-dependent manner in alcohol-naive and alcohol-dependent mice. Principal Component Analysis (PCA) revealed clear separation across genotypes and treatment conditions, indicating robust effects of neuronal *Fto* deletion and alcohol treatment (Fig.5a).

**Fig. 5.**
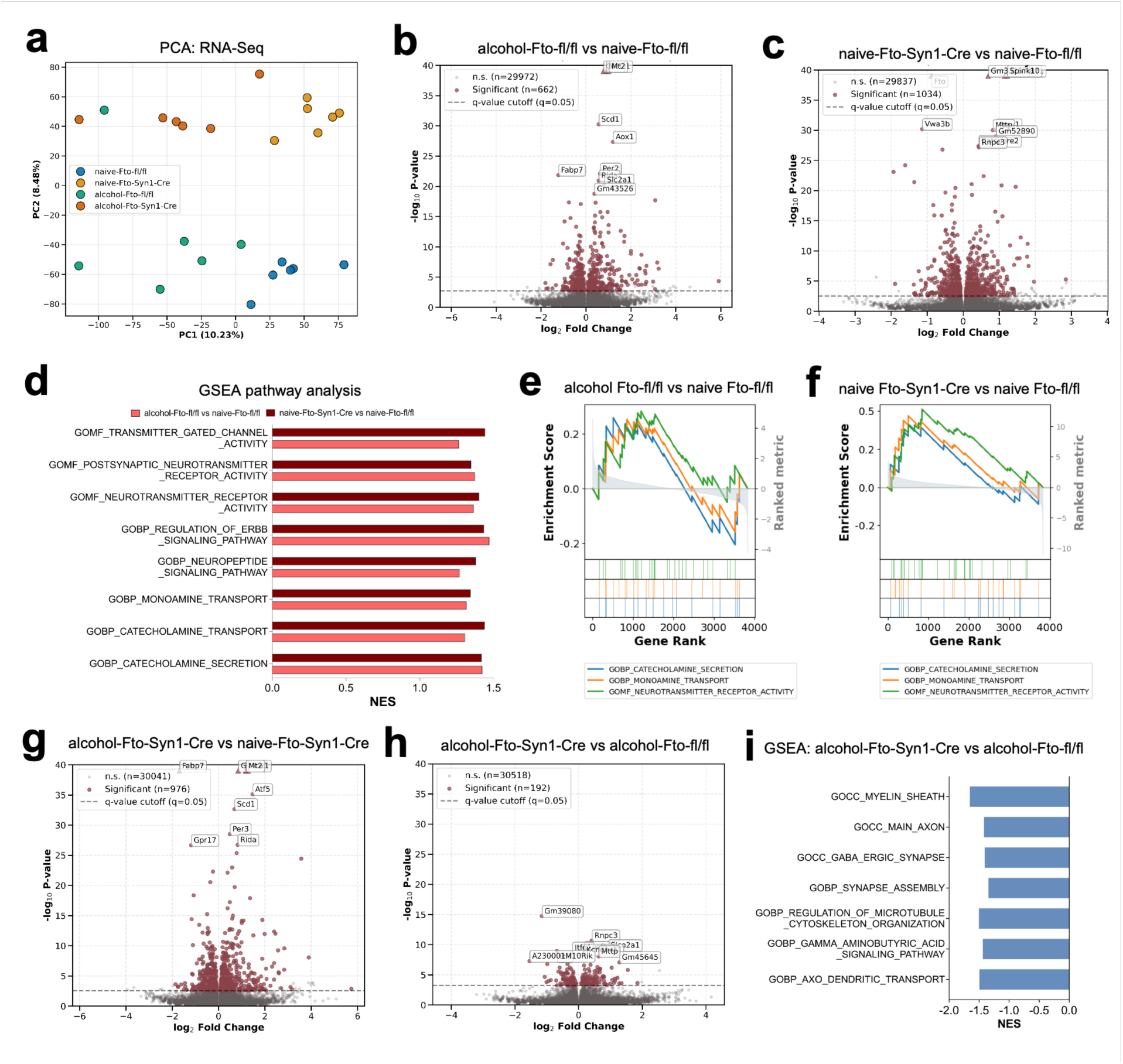
Alcohol dependence and neuronal *Fto* deletion reshape hippocampal gene expression and converge on neurotransmission pathways. a) Principal Component Analysis (PCA) of RNA expression detected by the RNA-Seq channel of the EpiPlex™ platform. Genotype and treatment replicates are indicated by color: alcohol-naive-*Fto-fl/fl* (blue, n=6), alcohol-dependent *Fto-fl/fl* (green, n=6), alcohol-naive *Fto-Syn1-Cre* (orange, n=6), alcohol-dependent *Fto-Syn1-Cre* (red, n=6). **b,c)** Volcano plots of differentially expressed transcripts in alcohol-dependent *Fto-fl/fl* (vs alcohol-naive *Fto-fl/fl*) (b) and in alcohol-naive *Fto-Syn1-Cre* (vs alcohol-naive *Fto-fl/fl* (c). **d**) NES (Normalized Enrichment Scores) of overlapping pathways in alcohol-dependent *Fto-fl/fl* (vs alcohol-naive *Fto-fl/fl*) and in alcohol-naive *Fto-Syn1-Cre* (vs alcohol-naive *Fto-fl/fl*). **e,f)** Representative enrichment plots of neurotransmission pathways from (d) commonly enriched in alcohol-dependent *Fto-fl/fl* (vs alcohol-naive *Fto-fl/fl*) (e) and in alcohol-naive *Fto-Syn1-Cre* (vs alcohol-naive *Fto-fl/fl*) (f). **g, h)** Volcano plots of differentially expressed transcripts in alcohol-dependent *Fto-Syn1-Cre* (vs alcohol-naive *Fto-Syn1-Cre*) (g) and in alcohol-dependent *Fto-Syn1-Cre* (vs alcohol-dependent *Fto-fl/fl* (h). **i)** NES of differentially enriched pathways in alcohol-dependent *Fto-Syn1-Cre* (vs alcohol-dependent *Fto-fl/fl)*. Transcripts with significantly increased and decreased expression were identified using DEseq2 (FDR-corrected P-value < 0.05). Selected pathways with P-value < 0.05 are shown.

A total of 662 (362 upregulated and 300 downregulated) differentially expressed genes were identified in alcohol-dependent *Fto-fl/fl* mice compared to their alcohol-naive littermates (Fig.5b). In addition, comparison between alcohol-naive *Fto-Syn1-Cre* and *alcohol-naive Fto-fl/fl* mice revealed 1034 (484 upregulated and 550 downregulated) differentially expressed genes in *Fto-Syn1-Cre* mice, with Fto among the most significantly downregulated transcripts (Fig.5c).

Importantly, GSEA revealed that alcohol exposure and neuronal *Fto* deletion converge at the transcriptional level. We compared pathway enrichment profiles derived from RNA-seq analyses between alcohol-dependent *Fto-fl/fl* mice (vs alcohol-naive *Fto-fl/fl*) and alcohol-naive *Fto-Syn1-Cre* mice (vs *alcohol-naive Fto-fl/fl*). Both conditions exhibited enrichment of overlapping pathways, particularly those related to neurotransmitter receptor activity, transmitter-gated channel activity, monoamine transport, catecholamine secretion, and neuropeptide and ERBB signaling (Fig.5d). Enrichment plots of representative pathways demonstrated similar patterns of gene enrichment across ranked gene lists in both comparisons (Fig.5e,f), indicating partially convergent transcriptional responses.

Notably, alcohol dependence induced a more pronounced transcriptional effect on *Fto-Syn1-Cre* than in *Fto-fl/fl* control mice. While 662 differently expressed genes were identified in alcohol-dependent *Fto-fl/fl* mice (Fig.5b), 976 (554 upregulated, 422 downregulated) were identified in alcohol-dependent *Fto-Syn1-Cre* in comparison to their alcohol-naive littermates (Fig.5g).

Finally, comparison between alcohol-dependent *Fto-Syn1-Cre* and alcohol-dependent *Fto-fl/fl* mice revealed 192 (100 upregulated, 92 downregulated) differently expressed genes, with *Fto* being one of the most significantly down-regulated genes in *Fto-Syn1-Cre* mice (Fig.5h). Pathways related to GABAergic neurotransmission were downregulated in alcohol-dependent *Fto-Syn1-Cre* mice in comparison to alcohol-dependent *Fto-fl/fl* (Fig.5i). Alcohol-dependent *Fto-Syn1-Cre* mice also showed a broad downregulation of neuronal structural and functional pathways (Fig.5i), potentially indicative of neurodegeneration-like changes.

Collectively, these findings demonstrate that neuronal *Fto* deletion amplifies and partially overlaps with alcohol-induced transcriptional remodeling in the hippocampus. All differentially expressed transcripts and pathways can be found in the in the Supplementary Tables (Supplementary Table 9, Supplementary Table 10, Supplementary Table 11, Supplementary Table 12, Supplementary Table 13, Supplementary Table 14, Supplementary Table 15, Supplementary Table 16). Quality control (QC) can be found in the Supplementary Information (Supplementary Figure 2).

## Discussion

Alcohol use disorder (AUD) is among the most consequential mental disorders worldwide and is a major contributor to global disease burden and premature mortality ^18,19^. Thus, there is a pressing need to better understand its complexity to identify new and more effective druggable targets ^20^. While canonical epigenetic gene regulation in the pathogenesis of AUD has been extensively explored ^21–23^, the role of internal RNA modifications or epitranscriptomic modifications remains uncharacterized. Neurons have especially high levels of m^6^A, which regulate key processes underlying synaptic plasticity, learning and memory formation ^12,13^. RNA m^6^A methylation is a reversible modification, and its removal is mediated by FTO ^11^, making it a key regulator of gene expression states ^9^. Here we provide the first evidence of a critical functional role of m^6^A methylation and its demethylase FTO in the effects of alcohol and in the vulnerability to develop AUD.

We generated mice with neuronal m^6^A hypermethylation through selective neuronal deletion of *Fto* and tested their alcohol sensitivity, alcohol intake and alcohol-dependence. Selective neuronal deletion of *Fto* resulted in heightened initial motivational response to alcohol in alcohol-naive *Fto-Syn1-Cre* mice. The transition from occasional use to addiction involves neuroplasticity and neuroadaptations that begin with changes in the mesolimbic dopamine system followed by broad neuroadaptations in other limbic circuits ^24^. *Fto* knockdown in dopaminergic neurons has been shown to potentiate cocaine-induced positive reinforcement and associative learning in conditioned place preference ^25^. Learning processes driven by positive or negative reinforcement are implicated in drinking trajectories as, for instance, early alcohol consumption is a predictor of subsequent drinking behavior in rodents ^26^. Unlike *Fto-fl/fl* controls, *Fto-Syn1-Cre* mice displayed no correlation between initial and dependent alcohol consumption, suggesting that *Fto* regulates how early reinforcing alcohol experiences are translated into persistent heavy drinking patterns. In humans, FTO genetic variants were suggested to increase susceptibility to obesity through altered sensitivity and learning from negative outcomes ^27^.

Our observation that *Fto-Syn1-Cre* mice required fewer alcohol exposures to achieve alcohol dependence-associated escalated drinking and exhibited enhanced relapse-like behavior suggests a predisposition to accelerated and heightened alcohol-induced neuroadaptive and plastic processes. Consistent with this, the GSEA of our epitranscriptomics and transcriptomics datasets revealed that *Fto* deficiency induced alterations in multiple signaling cascades known to govern synaptic plasticity, neurogenesis, and addiction-relevant neuroadaptations, such as MAPK-ERK signaling^28,29^, neuropeptide signaling ^30^, WNT-signaling ^31,32^ and Rho GTPase signaling ^33^.

Importantly, we also observed a partial convergence between alcohol-induced and *Fto*-dependent transcriptional remodeling in pathways governing neurotransmission and synaptic signaling. This overlap suggests that neuronal *Fto* deletion partially mimics core components of the alcohol-responsive transcriptional program in the hippocampus. Both conditions independently enriched pathways related to neurotransmission, transmitter-gated channel activity, monoamine transport, and synaptic signaling, indicating convergence on shared neuroplastic processes. These alterations might constitute the molecular basis of the increased vulnerability of mice with neuronal *Fto* deficiency. This model is consistent with the emerging view that m^6^A RNA methylation acts as a dynamic regulator of neuronal gene expression programs and is in keeping with previous reports implicating *Fto* and m^6^A in synaptic plasticity and memory formation ^12–14,34–36^.

We observed a critical potentiation of alcohol-induced locomotor responses in *Fto-Syn1-Cre* mice. Alcohol-induced hyperactivity is largely dopamine dependent ^37^. Increased locomotor response to cocaine was also observed in mice with *Fto* deficiency in dopaminergic neurons ^25^, as well as enhanced locomotor response to novelty ^38^. Notably, our RNA-Seq analysis revealed a significant upregulation of the *Dopa Decarboxylase (DDC)* gene, responsible for dopamine synthesis ^39^ and our GSEA pathway analysis revealed significant upregulation of catecholamine and monoamine transport and secretion pathways in alcohol-naive *Fto-Syn1-Cre* when compared to alcohol-naive *Fto-fl/fl*.

*Fto-Syn1-Cre* mice were also more susceptible to relapse to heavy drinking following a period of abstinence. Withdrawal from chronic alcohol decreases GABA neurotransmission ^40^. Insufficient GABAergic inhibition may result in withdrawal manifestations such as anxiety, hyperexcitability, insomnia, and seizures, which may contribute to alcohol-seeking behavior ^41^. GSEA analysis of our RNA-Seq dataset revealed a downregulation of GABAergic signaling pathways in alcohol-dependent *Fto-Syn1-Cre* mice compared to alcohol-dependent *Fto-fl/fl* mice during alcohol withdrawal, supporting an amplified and qualitatively distinct transcriptional response to alcohol in the absence of *Fto*.

Alterations in GABA dynamics are suggested also by the different phenotype of *Fto-Syn1-Cre* mice in the anxiety-related and sedative responses to acute alcohol. We observed that genetic inhibition of *Fto* markedly potentiates alcohol-induced anxiolysis and completely blunted anxiogenic responses. The subjective way in which individuals emotionally experience early alcohol exposures is a critical determinant for later AUD risk ^42–44^ and drinking to cope with anxiety or for tension reduction is consistently associated with greater risk of problematic use, underscoring the reinforcing impact of anxiolysis itself ^45,46^.

GABA receptors also mediate alcohol’s sedative and hypnotic properties ^16,17^. In agreement, we also report increased sensitivity to alcohol-induced hypnosis in *Fto-Syn1-Cre* mice, that could act as a limiting factor to excessive consumption, in keeping with the absence of significant effects of *Fto*-deficiency on alcohol intake during chronic heavy exposure. In support, *Fto* and m^6^A RNA methylation have been implicated in the regulation of stress response and anxiety ^47,48^ and *Fto* knockdown has been shown to increase the expression of GABAA receptor subunits ^49^, key molecular targets of anxiety manifestations of typical anxiolytic drugs, as well as alcohol ^16,50^.

Increased GABA_A_ receptor-mediated response to acute alcohol may underlie the enhanced alcohol-induced anxiolysis and sedative response that we observed in *Fto-Syn1-Cre* mice. GABA-hypersensitivity would also explain the exacerbation of the molecular signatures of alcohol-induced GABAergic withdrawal ^40,41,51,52^ that we observed in alcohol-dependent *Fto-Syn1-Cre* mice, as well as their propensity to relapse.

In summary, our findings identify RNA m^6^A methylation as a critical contributor to the maladaptive neuroplasticity that underlies excessive alcohol drinking and vulnerability to relapse that characterize alcohol addiction ^53^. The present behavioral, m^6^A-profiling and transcriptomic data suggest that RNA m^6^A methylation may increase individual vulnerability to AUD by predisposing neurons to alcohol-induced neuroadaptive changes that contribute to enhance the initial motivational response to alcohol, accelerate the onset of dependence and aggravate relapse behavior. The involvement of m^6^A dysregulation also in depression and stress responses, which are associated with AUD vulnerability ^54^ suggests that targeting m^6^A-regulated pathways may represent a promising direction for therapeutic intervention.

## Methods

### Animals

All experimental procedures were approved by the Institutional Animal Care and Use Committee of The Scripps research Institute and performed in accordance with the Association for Assessment and Accreditation of Laboratory Animal Care. *Fto^flox/flox^* mice were generously provided by Dr. Clemens (Johns Hopkins University). To generate brain neuronal specific *Fto* deletion, *Fto^flox/flox^* mice were crossed with *Syn1-Cre* mice (JAX 003966) to generate *Fto^+/flox^*/*Tg(Syn1-Cre)* mice, which were then crossed to *Fto^flox/flox^* mice to generate the experimental *Fto-fl/fl* and homozygous *Fto-Syn1-Cre*. Animals were housed in a climate-controlled vivarium on a 12 h/12 h reverse light/dark cycle (lights on 9:00 AM) and provided with free access to water and food (7012 Teklad LM-485 Rodent Diet, inotive, IN, USA) throughout the study.

### Behavioral procedures

#### Two-bottle choice (2BC) paradigm and Chronic intermittent access (CIE) to alcohol vapors

Male *Fto-fl/fl* and *Fto-Syn1-Cre* mice were singly housed and exposed to the conventional two-bottle alcohol [15% (v/v)] vs water choice (2BC) paradigm for 4 weeks (5 weekly 2h sessions) to measure initial alcohol intake in naive mice and obtain a baseline of alcohol-intake. Alcohol and water consumption were recorded by weighing the bottles before and after the drinking session. Subsequently, mice were exposed to intermittent ethanol vapor (CIE, 16h ON, 9h OFF) to induce escalated-dependent drinking. Statistically significant escalation was defined by p<0.05 vs the last week of baseline alcohol intake. Before being exposed to alcohol vapors, mice were injected IP with a solution of alcohol (1.5 g/kg, 15% w/v in saline) containing 68.1 mg/kg pyrazole and immediately placed into alcohol vapor chambers (La Jolla Alcohol Research, CA, USA) to reach a target Blood Alcohol Levels (BALs) of 150–200 mg/dL. Seventy-two hours following removal from the chambers, mice received access to water versus 15% v/v alcohol for 2 hours, and again over the next 4 days. Over the following weeks, mice were re-exposed to the alcohol vapor/control conditions and again tested for 2BC drinking for 5 days for a total of 4 alcohol vapor cycles each followed by 2BC. After the last cycle, mice were prevented from accessing alcohol for a total of 10 days to induce withdrawal and were then tested acutely with a 2BC test to measure reinstatement-like behavior. Mice were then re-exposed to a week of vapors and euthanized 24h after the last exposure. Body weight was taken every 4–6 days throughout the 2BC sessions and daily during the vapor exposure bouts. Food and water were available ad libitum, and mice were always group-housed except during the alcohol-drinking sessions. Two *Fto-fl/fl* and four *Fto-Syn1-Cre* died along the experimental procedure.

#### Elevated Plus Maze and Light and Dark Transition Test

The Elevated Plus Maze (EPM) procedures were conducted according to our previous studies ^55,56^. The EPM consisted of a central platform (5×5 cm, W×L) and two open arms (5×30 cm, W×L) aligned perpendicularly to two closed arms (5×30 cm, W×L) at the height of 40 cm from the ground. *Fto-fl/fl* and *Fto-Syn1-Cre* mice were administered with alcohol (1.5 g/kg) or saline and tested 5 minutes later. The spontaneous activity of mice was automatically recorded for 5 min. Data were calculated as the percentage of time spent in the open arms (*time spent in the open arms divided by the time spent in the open arms + time spent in the closed arms*) and as the percentage of open arm entries (*number of entries into the open arms divided by the number of entries into the open arms + number of entries into the closed arms*). The total number of entries was measured and reported as a measure of activity and to exclude sedation. The analysis was automatically performed by ANY-Maze software. Similarly, a separate cohort of *Fto-fl/fl* and *Fto-Syn1-Cre* mice was administered with alcohol (1.5 g/kg) or saline and tested at the Light and Dark Transition (LTD) test 5 minutes later. The spontaneous activity of mice was automatically recorded for 10 min. Data were calculated as the percentage of time spent in the light zone, as well as the percentage of entries in the light zone. The total locomotor activity was measured and reported. The analysis was automatic as the LTD arena was provided with infrared photo beams (Med Associates, USA).

#### Loss of Righting Reflex and alcohol metabolism

We assessed the effects of neuronal deletion of *Fto* on mice sensitivity to sedative/hypnotic effects of alcohol through the Loss Of Righting Reflex paradigm according to our previous studies ^57^. *Fto-fl/fl* and *Fto-Syn1-Cre* mice received an injection of alcohol 15% (w/v) [3.5 g/kg, IP]. Immediately after alcohol injection, when the mice showed the first signs of sedation, they were placed on their back on a V-shaped position into a cleaned cage, left undisturbed and monitored. Time to- and duration of-Loss Of Righting Reflex (LORR) were measured. In addition, blood samples were collected from the tip of the tail of each mouse when mice recovered the righting reflex (RORR) and after 2 hours from alcohol injection. BALs were measured as described above.

#### Blood alcohol levels (BALs)

BALs were determined by a conventional oxygen-rate alcohol analyzer (Analox Instruments, Atlanta, GA). The reaction is based on alcohol oxidation of alcohol oxidase in the presence of molecular oxygen (ethanol + O_2_ -> acetaldehyde + H_2_O_2_). The rate of oxygen consumption is directly proportional to the alcohol concentration. Blood was collected from tail tip, and BALs were collected along the CIE-2BC procedure and after the loss of righting reflex procedures.

#### Statistical analysis

Statistical analyses were performed using GraphPad Prism (version 11.0.0). Data distribution was assessed for normality using the Shapiro–Wilk test. Alcohol intake values during the first two-bottle choice (2BC) session did not meet normality assumptions and were therefore analyzed using a two-tailed Mann–Whitney test. Alcohol intake during baseline 2BC and chronic intermittent ethanol (CIE)-2BC sessions was analyzed using two-way repeated-measures ANOVA with genotype as the between-subjects factor and session as the within-subjects factor, followed by Tukey’s multiple comparisons test. Escalation of alcohol intake during dependence was assessed relative to the fourth baseline session (BL4). Alcohol intake during the withdrawal 2BC test was analyzed using a two-tailed Welch’s t-test. Pearson’s correlation analysis was used to evaluate the relationship between initial alcohol intake and subsequent drinking levels. Elevated plus maze (EPM) and light–dark test (LDT) behavioral parameters were analyzed using two-way ANOVA followed by Tukey’s multiple comparisons test. Time to loss of righting reflex (LORR) and time to recover the loss of righting reflex (RORR) were analyzed using a two-tailed Welch’s t test. Blood alcohol levels (BALs) measured during LORR procedures were analyzed using two-way repeated-measures ANOVA. Statistical significance was set at P < 0.05.

### EpiPlex™ m^6^A methylation encoding and RNAseq

After 24h of withdrawal, total RNA was extracted from dissected hippocampi of *Fto-fl/fl* and *Fto-Syn1-Cre* mice with RNeasy kit (Qiagen) and m^6^A methylation detection was performed using AlidaBio’s EpiPlex™ RNA Mod Encoding Kit (P/N 100124) using 1 µg purified total RNA isolated from mouse brain tissue as input. Sequencing libraries were prepared following kit instructions as previously written ^58^. In brief, RNA sample and spike-in controls were fragmented and ligated to an Illumina P7 adapter. Modified fragments were enriched on magnetic beads coated with modification-specific binders and corresponding Illumina P5 adapters, comprising a modification barcode (MBC) to capture and encode the RNA modification identity during reverse transcription, then PCR amplified using the EpiPlex™ Unique Dual Index Primer Kit (P/N 296001). A portion of the sample bypasses enrichment to provide a solution control library with RNA-Seq information. Ribosomal RNA was removed using the SEQuoia RiboDepletion Kit (Bio-Rad, P/N 17006487), and final libraries were prepared through additional PCR cycles. Paired-end libraries were sequenced to ∼50 - 80M reads on an Illumina NovaSeq X with the 10B (200 Cycle) Reagent Kit (P/N 20085595).

### EpiScout™ m^6^A peak calling and analysis

EpiPlex™ sequencing reads were processed using AlidaBio’s EpiScout™ v1.2 pipeline. The latest EpiScout^™^ software can be found in DockerHub (https://hub.docker.com/r/alidabio/episcout). Briefly, raw read quality was assessed using FastQC. Reads with length <30, mean quality <30, or NNN content >10% were discarded. Clean reads were then mapped to the mouse genome (GRCm39) and the EpiPlex™ internal spike-in control using STAR v2.7.11a ^59^. Genome mapped reads were split by MBC and deduplicated according to MBC and Unique Molecular Identifier (UMI) barcodes using samtools v1.20 ^60^ and custom Python and C++ libraries. Peak calling was performed using a Hidden Markov Model (HMM) based on the ratio of enriched and solution control read pileups, internal spike-in standard curve, and sequencing depth ratio between enrichment and solution control libraries. Peak quality was assessed using random forest model log-probabilities, represented as a PHRED scale q-score (random_forest_q). Peaks were annotated using the GRCm39 gtf file and a custom bedtools script ^61^.

### EpiScout™ differential m^6^A region identification analysis

Significant increases and decreases in m^6^A methylation were identified using the EpiScout™ pipeline. Briefly, peak intersection sets for each condition were identified using bedtools, requiring at least one replicate per condition to contain a peak that passes the random forest filter criteria. A count profile matrix from the union of peak regions between conditions was then generated, normalizing for library sizes, then extreme values using Trimmed Mean of M-values method ^62^. Briefly, differential m^6^A enrichment was assessed through the condition:sample interaction coefficient of a Poisson Generalized Linear Model, using an additional coefficient accounting for background transcript expression. p-values adjusted for multiple testing using the Benjamini-Hochberg false discovery rate procedure. Log₂ fold changes were calculated as the difference between normalized enrichment ratios across conditions. Separately, differential transcript expression analysis was performed on the EpiPlex^TM^ solution control library using DESeq2 ^63^. Significance for differential analysis was determined using adjusted P < 0.05.

### Gene Set Enrichment Analysis (GSEA) of m^6^A methylation and RNA-Seq expression profiles

Functional pathways relating to m^6^A methylation and expression were identified using GSEA ^64,65^. Briefly, count matrices for m^6^A and RNA-Seq profiles were provided as input to the GSEA command-line tool. GSEA was performed using mouse Molecular Signatures Database (MSigDB) gene sets from the Hallmark (MH), positional (M1), curated (M2), regulatory target (M3), Gene Ontology (M5), immunologic signatures (M7), and cell type signatures (M8) collections ^66^. Due to the large number of gene sets tested across collections, enrichment was evaluated using normalized enrichment scores (NES) and nominal P-values (P < 0.05), while FDR values are reported in Supplementary Tables.

## Acknowledgments

This work was supported by NIH Grant AA021667; IL was partially supported by training grant # T32 AA007456. Images were generated with BioRender.

## Author Contributions

RM, VRP and PPS conceived and designed the study. RM, IL, IT, RP and VRC performed the experimental procedures and collected the data. All authors contributed to the interpretation of the data. RM and PPS drafted the manuscript, and all authors

revised the manuscript critically for important intellectual content. All authors approved the final version of the manuscript for submission.

