## Supplementary Information for "N6-methyladenosine RNA methylation is a novel epitranscriptomic regulator of excessive alcohol drinking and vulnerability to relapse"

### Blood alcohol levels along 2BC-CIE procedures

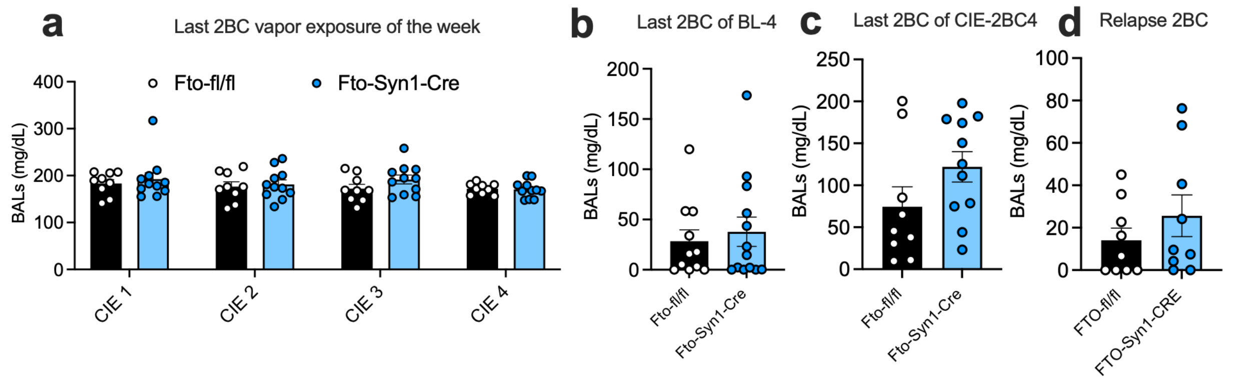


**Supplementary Figure S1.** Blood was collected from the tail tip after the last weekly session of CIE **(a)**, after the last 2BC session of basal drinking **(b)**, after the last 2BC session of the last week of CIE-2BC **(c)** and after the acute 2BC performed in withdrawal **(d)**. No significant effects were detected.

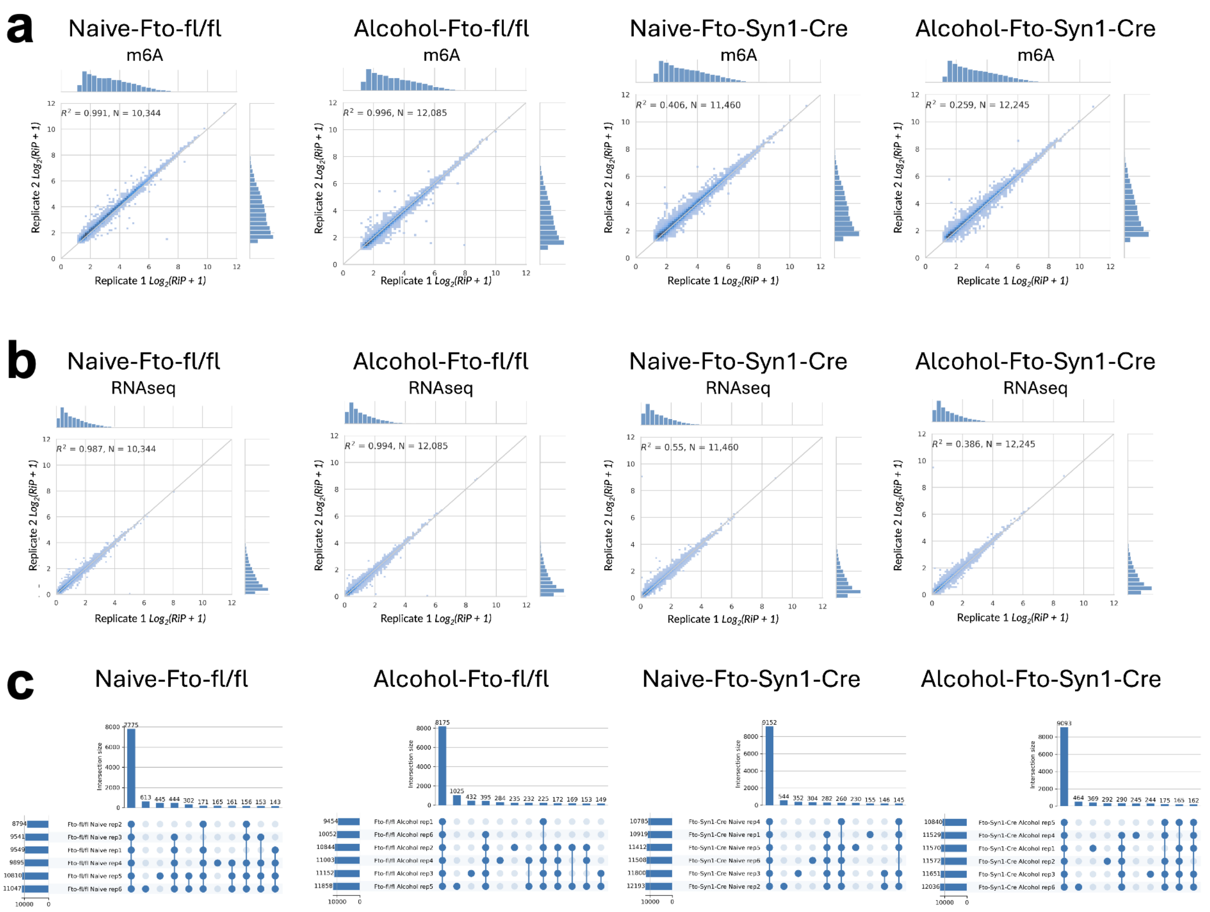


**Supplementary Figure 2. Quality control figures for m$^6$A modification detection by EpiPlex™. a)** Correlation plots of m$^6$A scaled Reads in Peaks (RiP) and **b)** Scaled Solution Control counts (RNAseq counts). Six replicates of the four treatment conditions are shown. The average scaled RiP of three replicates are compared against the other three. The R-squared value of N intersecting peaks is shown. **c)** Upset plots depicting unique sets of m$^6$A peaks belonging to combinations of replicates. Histograms on the left side of the plot show the total number of peaks per replicate, and bar plots show the total number of peaks per set.
